# Estimation of microtubule-generated forces using a DNA origami nanospring

**DOI:** 10.1101/2022.04.19.488748

**Authors:** Ali Nick Maleki, Pim J. Huis in’t Veld, Anna Akhmanova, Marileen Dogterom, Vladimir A. Volkov

## Abstract

Microtubules are dynamic cytoskeletal filaments that can generate forces when polymerizing and depolymerizing. Proteins that follow growing or shortening microtubule ends and couple forces to cargo movement are important for a wide range of cellular processes. Quantifying these forces and the composition of protein complexes at dynamic microtubule ends is challenging and requires sophisticated instrumentation. Here we present an experimental approach to estimate microtubule-generated forces through the extension of a fluorescent spring-shaped DNA origami molecule. Optical readout of the spring extension enables recording of force production simultaneously with single-molecule fluorescence of proteins getting recruited to the site of force generation. DNA nanosprings enable multiplexing of force measurements and only require a fluorescence microscope and basic laboratory equipment. We validate the performance of DNA nanosprings against results obtained using optical trapping. Finally, we demonstrate the use of the nanospring to study proteins that couple microtubule growth and shortening to force generation.

## Introduction

Microtubules are dynamic polymers that can exert pushing and pulling forces when they grow and shorten. Microtubule-generated forces are important at various stages of the cell cycle in a variety of cell types and contexts (reviewed in Gudimchuk and McIntosh, 2021). One of the most well-studied processes relying on microtubule-generated forces is mitotic cell division, when the ends of microtubules pull on the centromeric regions of chromosomes through protein structures called kinetochores (Musacchio and Desai, 2017). The microtubule-kinetochore interface is force-sensitive: tension at the centromere is thought to be converted into a biochemical signal that silences the mitotic checkpoint. However, the nature of the force-sensor that responds to the microtubule-generated tension in the kinetochore is still unclear (reviewed in Audett and Maresca, 2020).

Precise measurements of microtubule-generated forces and responses to these forces are challenging *in vivo*, because of crowded cellular environments, a multitude of differently directed forces that are exerted by different cellular components, and difficulties in incorporating force-measuring equipment into the cell. The method of choice for precision force measurements has been *in vitro* reconstitution and optical trapping. An optical trap holds a plastic or glass sphere (bead) in the centre of a tightly focused infrared laser beam. Bead displacement from the beam can be monitored with nanometre precision using sophisticated optical equipment (Baclayon et al., 2017; Nicholas et al., 2014). Forces measured using beads coated with microtubule-binding proteins or purified kinetochore particles provided important insights into the action of a microtubule as a motor (Grishchuk et al., 2005), the force-dependent stabilization of kinetochore-microtubule interface (Akiyoshi et al., 2010; Miller et al., 2016), and molecular determinants of kinetochore-mediated stabilization of microtubule ends (Huis in ‘t Veld et al., 2019; Volkov et al., 2018).

While having outstanding force-and time-resolution, the optical trapping approaches present several challenges. First, building an optical trap requires optical and engineering expertise, and commercial systems are expensive. Second, the trap acts on the centre of a bead, while force-generating biomolecules act on the surface of the bead, which can create asymmetry of the applied force in the bead-microtubule system (Pyrpassopoulos et al., 2020; Volkov et al., 2013). Finally, to study effects of force on dynamics of microtubule-binding or kinetochore proteins, one has to record these dynamics simultaneously with force. Addition of single-molecule fluorescence imaging to an optical trap is technically demanding (Deng and Asbury, 2017; Lang et al., 2004; Lee et al., 2007) and not widely accessible.

Here we present a method to simultaneously measure microtubule-generated forces and single-molecule fluorescence intensities *in vitro* without an optical trap. The method is based on a previously described DNA origami spring-shaped structure, designed to determine forces directly from its extension (Iwaki et al., 2016). The nanospring is assembled by folding a long single-stranded DNA with short DNA oligos (staples), resulting in a spring-shaped bundle of four DNA strands (Figure 1A, (Iwaki et al., 2016)). We provide detailed instructions to modify the original spring design, purify the DNA nanosprings, and use them as sensors for forces produced by growing and shortening microtubules.

**Figure 1.**
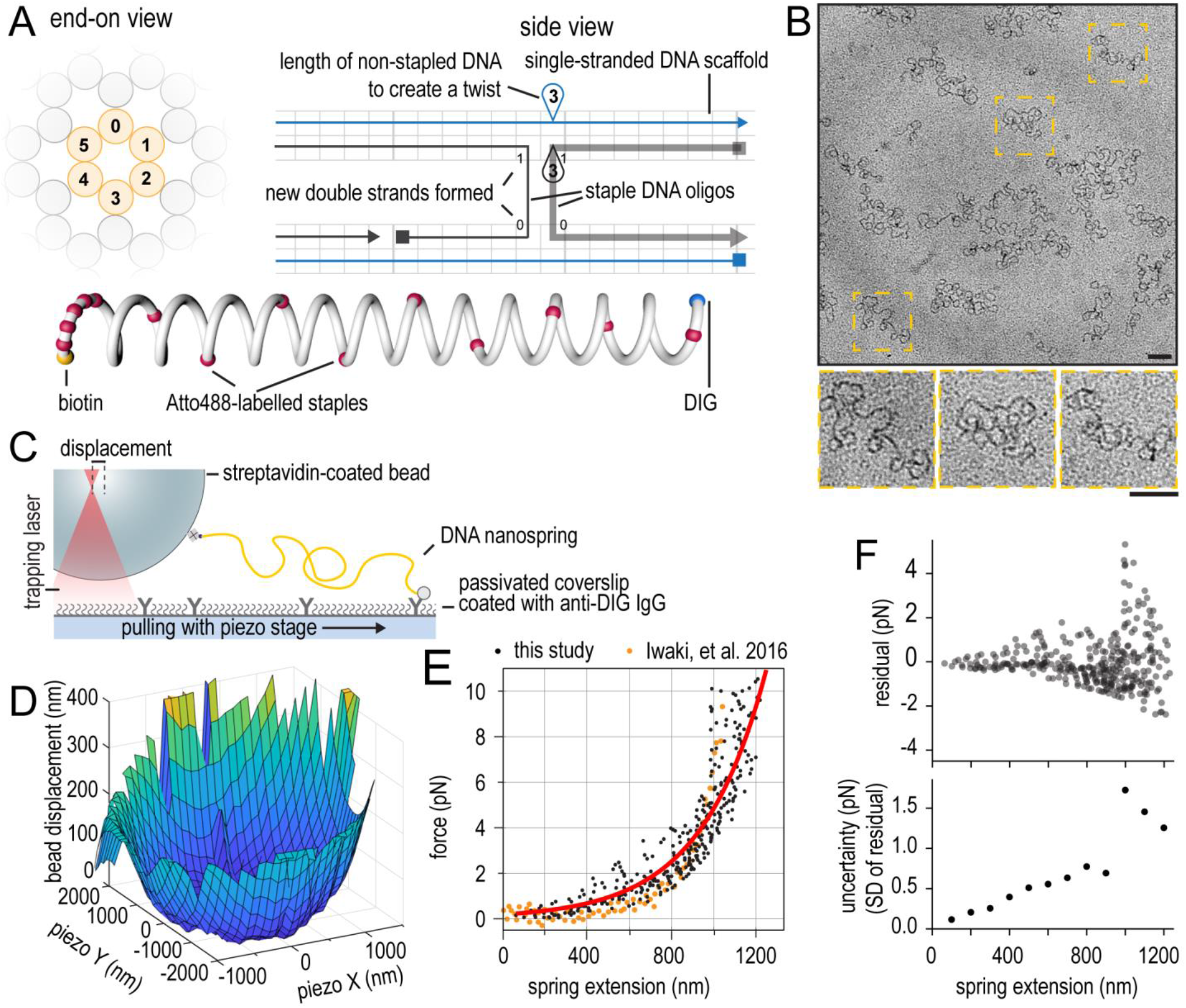
Design and calibration of the DNA origami nanospring. (A) An example of a slice panel (left) and a path panel (right) of the DNA origami design in caDNAno. Below: a schematic illustration of the nanospring with color-coded staples (yellow: biotin, red: fluorescently labelled staples, blue: DIG). (B) Transmission electron microscopy of nanosprings negatively stained by 2% uranyl acetate. Insets are zoomed in pictures of the marked regions. Scale bars: 100 nm. (C) Schematic diagram of an experiment with surface-bound nanospring being stretched using a bead in an optical trap. (D) A typical result showing nanospring extension (vertical axis) as a function of the 2D coordinates of the microscopic stage (horizontal axes X and Y). The bell-like shape of the curve signifies a single attachment point for the nanospring. (E) Force-extension curve resulting from measurements using six nanospring-attached beads (black), overlayed on top of the results presented in the original publication (orange, (Iwaki et al., 2016)). Solid red line shows results of exponential fitting. (F) Residuals of exponential fitting presented in (E) (top), and standard deviation of residuals binned by spring extension (bottom).

We provide typical results in three different *in vitro* systems. First, we validate the nanospring-based estimation of force against optical trapping by measuring the stall force of dynein motors walking along the stabilized microtubules. We then demonstrate the use of the nanosprings to measure the forces generated by dynamic microtubules. Focusing on forces produced by microtubule growth, we reconstitute the forces exerted through an EB3 comet following the growing microtubule ends and pulling on nanospring-bound cargo containing an EB-binding SxIP motif. Finally, we focus on forces produced by shortening microtubule ends in the context of kinetochore-microtubule interactions. We attach human kinetochore complex Ndc80 to the nanospring, and monitor spring extension simultaneously with the binding and unbinding of the Ska complex, another microtubule-binding kinetochore component. Using the DNA origami nanospring, we demonstrate that the presence of additional copies of Ndc80, but not Ska, increases the amount of force that shortening microtubule ends transmit to their cargo.

## Results

### Design and calibration of the nanosprings

To make the DNA nanospring, we adapted the reported DNA origami design (Iwaki et al., 2016) to other single-stranded DNA scaffolds (see Materials and Methods for a detailed protocol). The spring was assembled into a four-stranded DNA bundle with a three nucleotide-long hairpin inserted at every 14 bp, creating an offset to twist the spring (Figure 1A). DIG-and biotin-labelled DNA staples were introduced at the termini of the spring for attachment to the molecules of interest. Further, nine Atto488-labeled staple oligos were evenly distributed along the length of the spring, and another five Atto488-labeled oligos added next to the biotin-labelled end (Figure 1A). We validated the purity and folding of our DNA nanosprings using negative stain electron microscopy (EM) (Figure 1B). To calibrate the force-extension profile of the springs, we attached the springs to the glass surface of a flow chamber using anti-DIG IgG, and used 1 μm streptavidin-coated beads bound to the springs via biotin-streptavidin linkage (Figure 1C). Upon trapping a bead that was bound to the coverslip via the nanospring, displacement of the bead from the trap was measured as the flow chamber was moved using a piezo-driven stage in 100 nm steps following a 2D matrix (Figure 1D). After radial averaging of the force-extension data for 6 beads and accounting for the bead radius, the resulting force-extension curve was identical to the previously published one (Iwaki et al., 2016). Fitting the data to an exponential growth equation produced the following relationship which we used further to convert nanospring extension into force (Figure 1E):

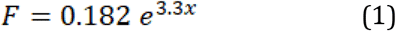

where F is force in piconewtons and x is nanospring extension in micrometres. Using fit residuals to estimate the uncertainty of force measurement from spring extension, we found that force can be determined within ± 1 pN for nanospring extensions below 1000 nm (Figure 1F).

### Validation of nanospring-based force measurements

To benchmark the performance of the nanospring in measuring single-molecule forces, we used biotinylated dynein motor domains from S. cerevisiae (Baclayon et al., 2017; Reck-Peterson etal., 2006). In this experiment, the nanospring was attached to anti-DIG IgG on the coverslip via DIG, and a streptavidin-coated Qdot bound both biotinylated dynein and a biotinylated DNA staple at the end of the nanospring (Figure 2A). We used total internal reflection fluorescence (TIRF) microscopy to record images of microtubules, Atto488-labelled nanosprings, and Qdot565-bound dynein at the end of the nanospring. Due to dimensions of the nanospring spanning several hundreds of nanometres, the best contrast was achieved using imaging mode with deeper penetration than TIRF, such as HiLo (Tokunaga et al., 2008), or intermediate settings between TIRF and epifluorescence. Using these settings, we could readily observe spring-bound Qdots walking along the microtubule, extending the springs, and stalling upon reaching the stall force of dynein (Figure 2B). To determine spring extensions from kymographs such as the one presented in Figure 2B, we used two methods. First, to determine the length of the spring, in each line of a kymograph containing the Atto488 signal, we measured the centre position of pixels that were brighter than the background fluorescence level. Second, to determine the position of the end of the spring, we fit a gaussian to each of the lines of a kymograph containing the Qdot-565 signal. Both of these measurements yielded estimates of spring extension (Figure 2C). However, subpixel localization of the Qdot position provided less noisy data (Figure 2D) and was therefore used in further analysis to determine the position of the spring end. Nanospring length was then converted into stalling force using Eq 1.

**Figure 2.**
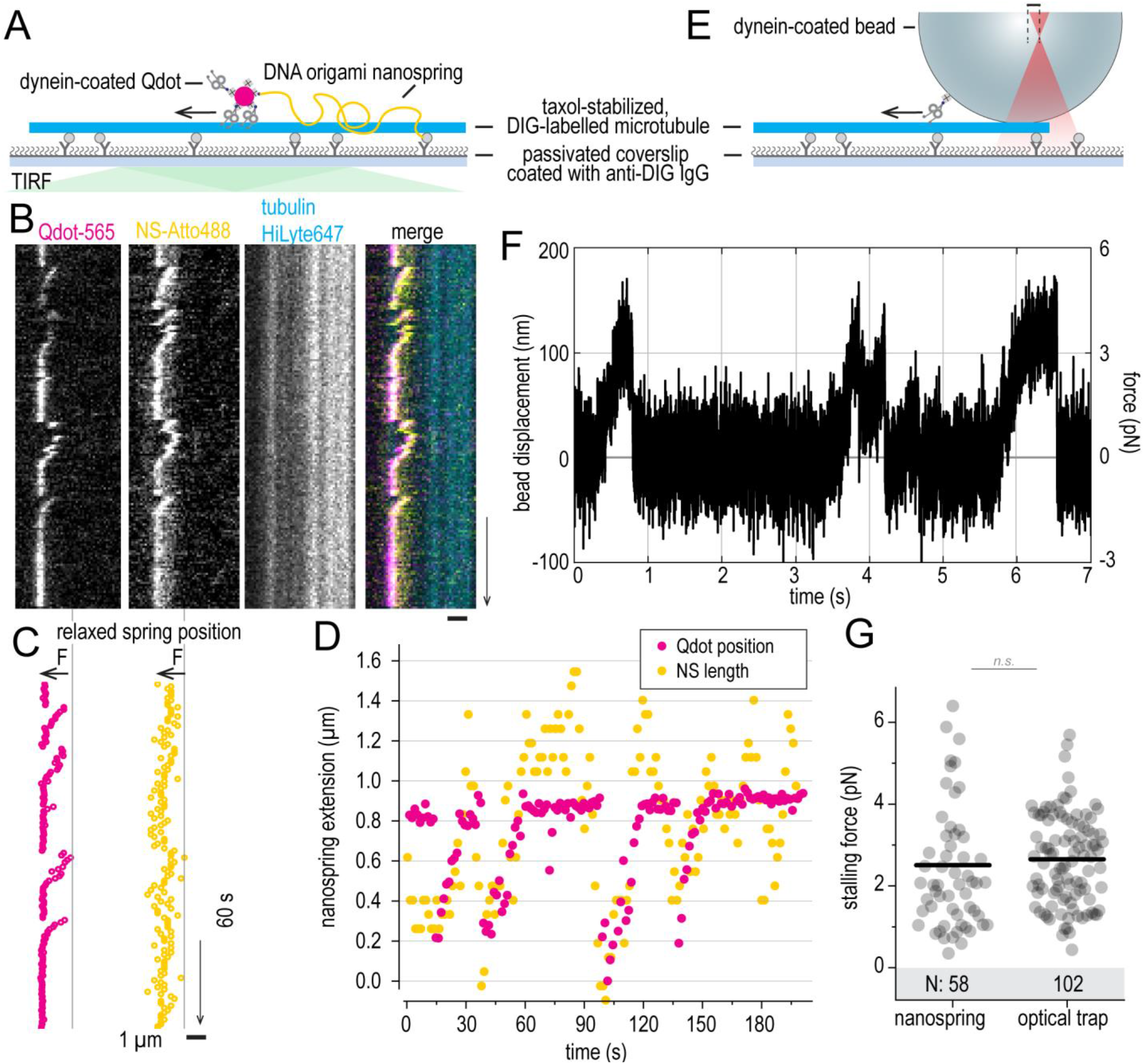
Analysis of spring extension and validation of force measurements. Experiment schematics (A) and a typical kymograph (B) showing coverslip-bound taxol-stabilized microtubule (cyan), Atto488-labelled nanospring (yellow) and dynein-conjugated Qdot-565 (magenta). (C) Coordinates on the nanospring end (Qdot, left), and nanospring middle (right) obtained by analysing the kymographs in B using the Julia script (see Materials and Methods for details). Vertical line shows the coordinates of the relaxed spring, arrows show direction of dynein-generated force. (D) Nanospring extension measured using these two methods and plotted as a function of time. (E) Experimental setup to estimate dynein stall force using an optical trap. (F) Typical time trace of a dynein walking against the applied force, recorded in an optical trap. (G) Dynein stalling forces measured using nanospring (left) and optical trapping (right). Circles: individual stall events, line: median, number of measurements is shown in the shaded area. Scale bars: horizontal (1 μm), vertical (60s).

As a control, we used optical trapping to measure the stall forces of bead-bound dynein (Figure 2E). Optical trapping provided high temporal and spatial resolution (Figure 2F), however the stall force values extracted from both force measurement methods were similar (Figure 2G). These results are consistent with other reports of *S. cerevisiae* dynein stall force (Gennerich et al., 2007; Laan et al., 2012).

### Estimation of forces generated by growing microtubules

Growing microtubule ends recruit end-binding (EB) proteins in the shape of a comet; these comets in turn recruit a number of secondary proteins that carry EB-binding SxIP motifs (Honnappa et al., 2009). The affinity of an EB comet to SxIP-containing proteins was reported to generate sub-pN forces that could extend membranes, transport actin filaments along with microtubule growth, and reverse the direction of a kinesin-14 motor (Alkemade et al., 2022; Molodtsov et al., 2016; Rodríguez-García et al., 2020). While prior measurements were performed using optical trapping, measuring sub-pN forces using this method is challenging, because it is easy to lose a bead from a soft trap. We therefore thought that the DNA nanospring, with its high precision in low-force regime could provide an advantage (Figure 1E). To couple nanospring extension to microtubule growth, we used a nanospring-bound Qdot705 coated with mCherry-tagged and biotinylated C-terminal fragment of human MACF2 (Rodríguez-García et al., 2020) in the presence of dynamic microtubules and EB3 (Figure 3A). In these conditions we observed nanosprings getting extended in the direction of microtubule growth when interacting with the EB3 comets through MACF (Figure 3B). This experiment highlights how protein complexes at the interface of the nanospring and the microtubule end can be directly visualized using fluorescence microscopy. Our results are consistent with observations using optical trapping, where a MACF-coated bead was interacting with a growing microtubule end (Figure 3C). In both conditions, we observed forces in sub-pN range lasting for many seconds (Figure 3B,D). On average, nanospring-measured forces were smaller than optical trap-measured ones (Figure 3E). This difference could be related to different amount of MACF molecules interacting with a comet in each case: no more than 20 in case of Qdot-nanospring, and several thousand in case of MACF-coated beads (Rodríguez-García et al., 2020).

**Figure 3.**
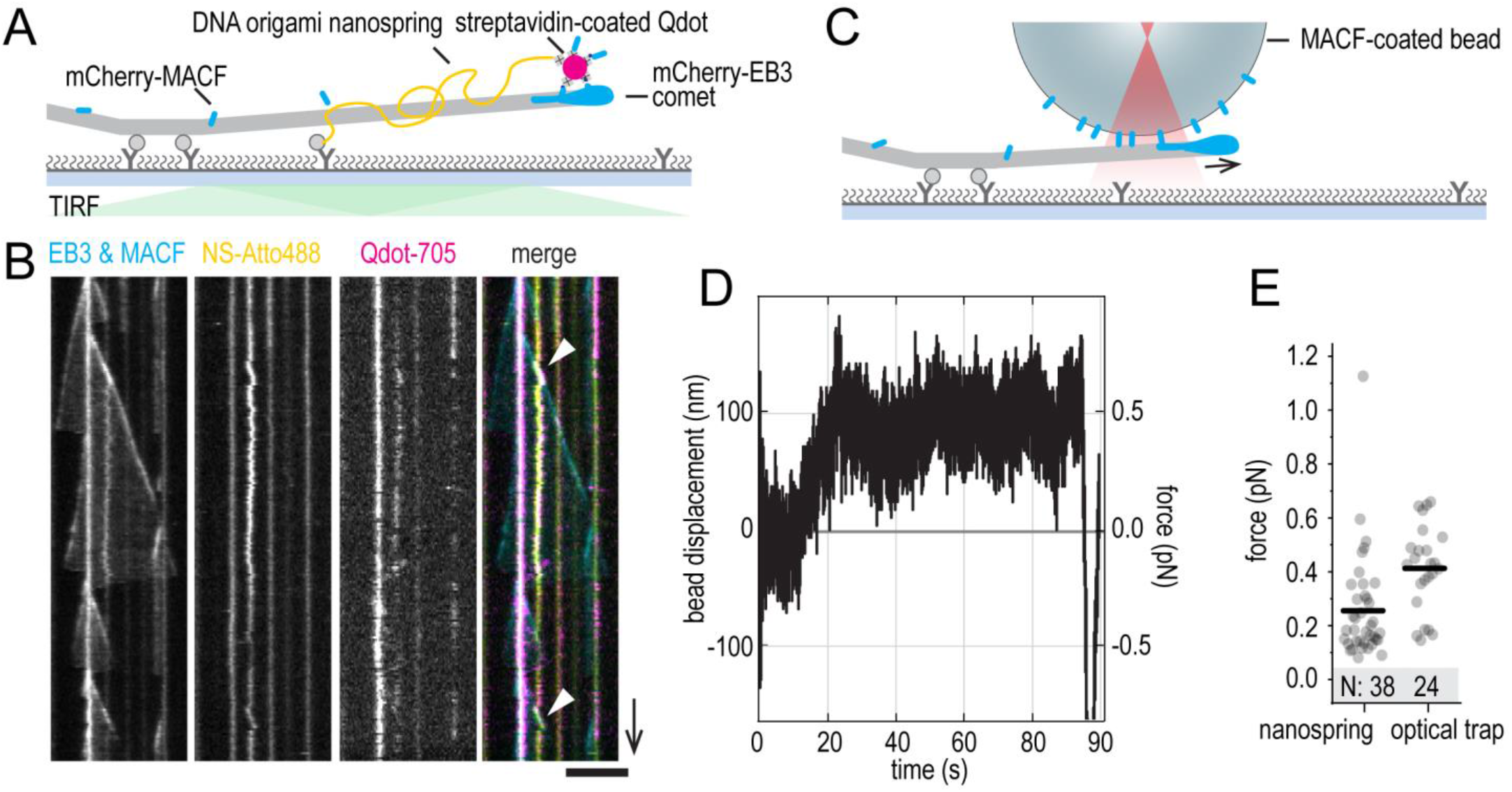
Estimation of microtubule growth force using DNA origami nanospring. Experimental setup (A) and typical kymogrpahs (B): microtubules are grown from coverlip-attached GMPCPP-stabilized seeds (grey). Yellow shows the nanospring. Streptavidin-coated Qdot-705 (magenta) binds to the nanospring and is further saturated using mCherry-tagged MACF C-terminus (cyan) (see Materials and Methods for details). MACF is concentrated at the growing microtubule end thanks to the mCherry-tagged EB3 also present in solution (cyan). Arrowheads show events of nanospring being stretched by growing microtubule ends. (B) Events of growing microtubule ends pulling on the nanosprings via MACF are shown using arrowheads. (C) Experimental setup to estimate the MACF-transmitted force using the optical trap. (D) Typical time trace of a glass bead following microtubule growth against the applied force. (E) Forces measured in the direction of microtubule growth using MACF-conjugated nanosprings or MACF-coated beads in an optical trap. Scale bars: horizontal (5 μm), vertical (60s).

### Estimation of forces generated by microtubule shortening

We have previously shown that multimerization of the human kinetochore complex Ndc80 enables it to follow microtubule shortening against an applied force (Volkov et al., 2018). Multiple copies of Ndc80 oligomers could stall microtubule shortening, duration of these stalls was increased in presence of another multi-protein kinetochore complex known as Ska, which cross-linked Ndc80 and microtubules (Huis in ‘t Veld et al., 2019). However, these observations were performed using bead-bound Ndc80 in an optical trap, in conditions preventing us from having precise information about the number of Ndc80 and Ska copies interacting with the force-generating microtubule ends.

To study the Ndc80:Ska:microtubule end-tracking system in single-molecule conditions, we used streptavidin-oligomerized Ndc80 bound to biotinylated nanosprings in presence of dynamic microtubules and Ska (Figure 4A). Using TIRF microscopy, we could simultaneously record microtubule dynamics, position of the spring-bound Ndc80, and dynamics of Ska binding to both microtubules and Ndc80 (Figure 4B). Forces measured using nanosprings carrying a single Ndc80 trimer in absence of Ska were similar to previously reported forces measured using beads sparsely coated with the Ndc80 trimers in an optical trap (Figure 4C) (Volkov et al., 2018). We then compared these forces to nanospring-measured forces recorded when Ska signals colocalized with the spring-bound Ndc80 (Figure 4B, top). As reported previously, at low Ska concentrations the Cdk1 phosphorylation of SKA3 C-terminus enhanced Ska:Ndc80 interactions (Huis in ‘t Veld et al., 2019), which is evident from the higher frequency of Ska-positive force events (Figure 4C). At 100 nM Ska, Cdk1 phosphorylation was no longer necessary for Ska:Ndc80 binding, however we did not observe a difference in nanospring-measured force resulting from the presence or absence of Ska during force development (Figure 4C).

**Figure 4.**
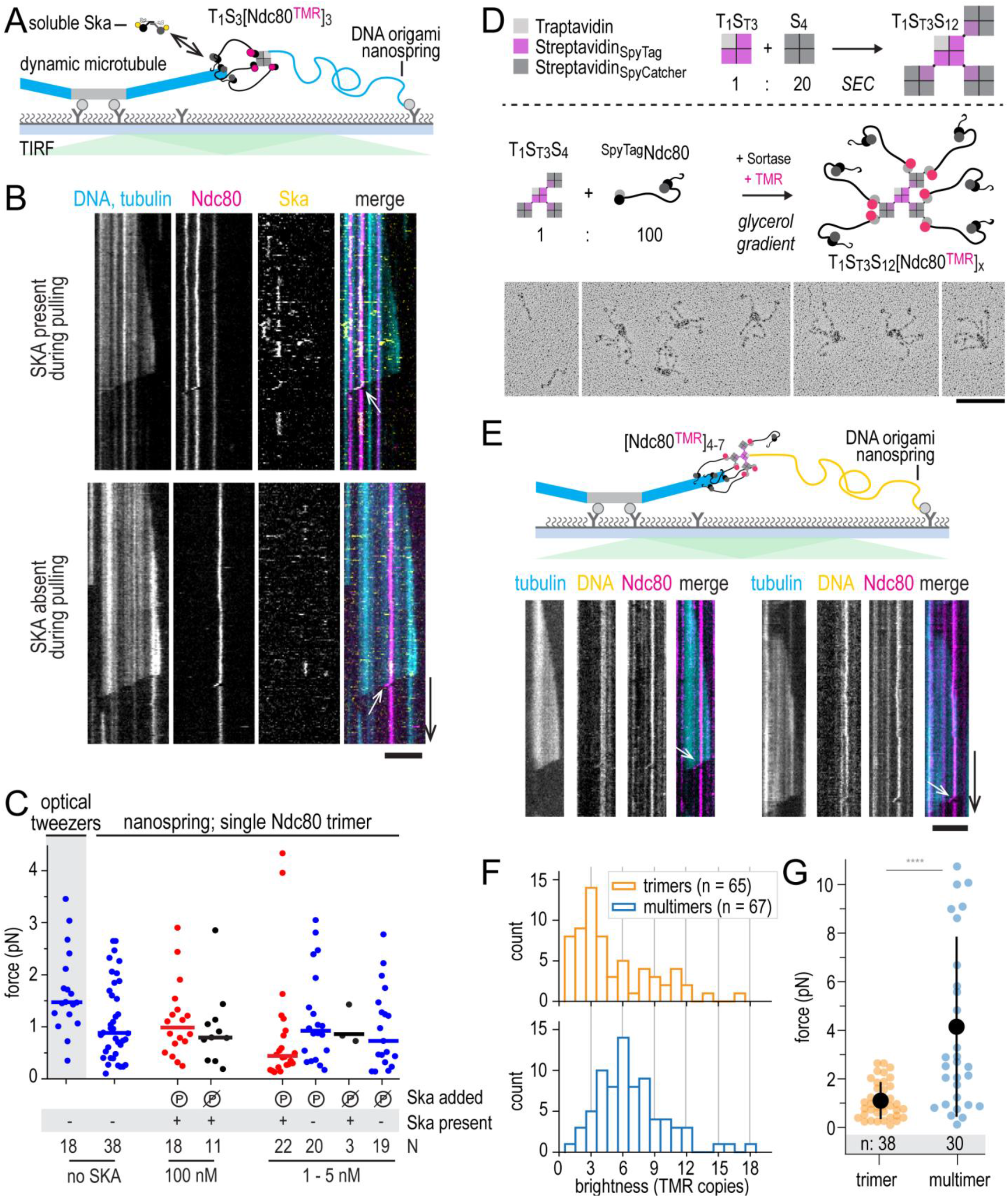
Estimation of microtubule shortening force using DNA origami nanospring. (A) Experimental setup: GMPCPP-stabilized microtubule seeds (grey) are attached to the coverslip and nucleate dynamic fluorescently labelled microtubules (cyan). Atto488-labelled nanospring (cyan) is conjugated to Ndc80-TMR trimers (magenta) oligomerized using streptavidin. Ska complex labelled with HiLyte-647 (yellow) is present in solution and can bind both to Ndc80 and to microtubules. (B) Typical results showing shortening microtubule ends pulling on nanospring-conjugated Ndc80 trimers (arrows). Top example: Ska is bound to the Ndc80 trimer during force development. Bottom example: Ska is present in the sample, but not bound to the Ndc80 trimer during force development. (C) Forces measured using a single Ndc80 trimer following microtubule shortening against the force exerted by the stretched nanospring. Shaded area: stalling forces measured using beads sparsely coated with Ndc80 trimers in an optical trap (Volkov et al., 2018). Circles: individual measurements, lines: median. Cdk1 phosphorylation of SKA3 leads to increased recruitment of Ska to Ndc80 trimers during force development (Huis in ‘t Veld et al., 2019), but has no effect on the measured force within the Ska concentrations in the range between 1 and 100 nM. (D) T1S_T_3S_C_12 assemblies were generated from T1S_T_3 and S_C_12 tetramers, purified using size-exclusion chromatography, and incubated with Ndc80. See Supplementary Figure S2 for more information. Multimeric Ndc80 assemblies were purified on a density gradient and inspected by electron microscopy after low-angle platinum shadowing. Monomeric Ndc80 complexes are shown in the first box, the next boxes contain representative micrographs with multimeric Ndc80 modules. (E) Experimental setup and typical kymographs showing DNA nanosprings (yellow) extended by Ndc80 multimers (magenta) following the shortening ends of dynamic microtubules (cyan). (F) Brightness of Ndc80 trimers (orange) and multimers (blue) expressed as a copy number of Ndc80-TMR. (G) Forces calculated from nanospring extension for a single Ndc80 trimer (orange) and a single Ndc80 multimer (blue). Scale bars: horizontal (5 μm – B,E; 100 nM – D), vertical (60s).

The failure of Ska to increase the Ndc80-mediated force was consistent with our earlier estimates using optical trapping, where the presence of Ska mainly affected the duration of stalls, and to a lesser extent the stalling force (Huis in ‘t Veld et al., 2019). We wondered however if there were conditions in which we could capture higher microtubule-generated forces using nanospring-bound Ndc80. One way to increase the amount of transmitted force is to engage more copies of a force-coupling molecule (Volkov, 2020; Volkov et al., 2018). To this end, we generated objects containing four modified streptavidin tetramers which could bind up to nine Ndc80 copies in total (see Materials and Methods and Supplementary Figure S2 for details). Examining the resulting Ndc80 multimers by electron microscopy, we found Ndc80 multimers with varying stoichiometries. Typical Ndc80 multimers contained 4-7 Ndc80 arms (Figure 4D). Attaching these multivalent objects to the biotinylated nanosprings, we could measure the force they transmitted from the microtubule shortening, and their brightness (Figure 4E). Consistent with our observation by EM, the distribution of Ndc80 copy number was quite wide, peaking at 6 Ndc80 copies per object (Figure 4F). Consistently with increased Ndc80 copy numbers, Ndc80 multimers transmitted up to 10 pN of microtubule-generated force, compared to a maximum of 3 pN transmitted by a single Ndc80 trimer (Figure 4G). Contrary to the experiments with bead-bound Ndc80 trimers, we did not observe force-dependent microtubule rescue with Ndc8 multimers bound to nanosprings despite forces in the 5-10 pN range.

## Discussion

Classically, force-production by the mitotic spindle and force-sensitivity of the mitotic checkpoint were studied using microneedles inserted into a dividing cell (Li and Nicklas, 1995; Nicklas, 1983). Recent developments of this approach yielded important insights into the organization of mitotic spindle (Long et al., 2020; Suresh et al., 2020), but precise quantification of forces *in vivo* remains challenging. Alternatively, tension at the centromeric region of the chromosomes could be estimated indirectly, based on assumptions about stiffness of stretchable elements in the cells (Harasymiw et al., 2019; Mukherjee et al., 2019).

The third method of estimating microtubule-generated tension *in vivo* is based on Förster resonance energy transfer (FRET) sensors. With a FRET sensor, the efficiency of energy transfer between donor and acceptor fluorophores depends on the distance between these fluorophores, allowing reading out tension using fluorescence intensity of the acceptor fluorophore (Cost et al., 2015; Grashoff et al., 2010). Although the FRET approach provided important evidence regarding the role of tension in regulating the kinetochore-microtubule attachments *in vivo*, multiple copies of microtubule-binders interacting with multiple microtubule ends result in ensemble readouts that are challenging to interpret at single-molecule level (Kuhn and Dumont, 2019; Liu et al., 2009; Suzuki et al., 2016; Ye et al., 2016; Yoo et al., 2018).

The DNA nanospring is a force-sensor that outputs a force signal directly through measurement of its extension. Potential advantages of nanosprings include multiplexed force measurements in a single microscopic field of view, simultaneous visualization of additional factors being recruited during force production (Figure 4B), and implementation in laboratories without access to an optical trap. Compared to an optical trap, nanosprings have a lower time resolution, limited to 1-100 Hz by the frame rate of the timelapse image acquisition as opposed to kHz range in an optical trap. At the same time, the use of the nanospring alleviates concerns related to the ‘lever arm’ effects arising from the geometry of microtubule-bead connection (Pyrpassopoulos et al., 2020; Volkov et al., 2013).

Because of a non-linear force-extension curve (Iwaki et al., 2016), the nanospring provides particularly clear readout in the sub-pN force range. This enabled us to measure forces generated by a protein complex tracking growing microtubule ends (Figure 3). Although the same phenomenon could also be observed using optical trapping (Alkemade et al., 2021; Molodtsov et al., 2016; Rodríguez-García et al., 2020), measurement of such small forces is usually technically challenging. It should be noted that nanosprings are less precise than optical tweezers in force estimation (uncertainty ~ 0.2 - 1.0 pN, Figure 1F, compared to ~ 0.1 pN or lower for optical tweezers).

Forces generated by shortening microtubules can sometimes exceed 10 pN (Akiyoshi et al., 2010; Volkov et al., 2013), the upper limit of the nanospring’s force-extension curve (Figure 2E). At higher forces the DNA origami structure might unfold (Engel et al., 2018) or detach from its surface anchor (Neuert et al., 2006). A possible improvement of the attachment strength is DIG10.3 instead of anti-DIG IgG (Tinberg et al., 2013; Van Patten et al., 2018). In addition, multiple DIG-and biotin-labelled staples could be introduced to share the load evenly. In our experience, the DNA origami nanosprings were stable in microtubule force experiments for up to 30-60 minutes. To further improve the lifetime of the nanospring under force, ligation (Ramakrishnan et al., 2019) or chemical crosslinking of strands (Rajendran et al., 2011; Thomas et al., 2022) after purification could be beneficial.

In this study, nanosprings provided similar force amplitude values compared to published optical trapping experiments in different experimental settings. MACF-mediated force transmission resulted in consistently lower forces produced by microtubule growth and measured using the nanospring. We hypothesise that the nanospring could be particularly sensitive in sub-pN regime, where extension of the spring within the 200-300 nm range leads to estimation of the force with 0.2-0.3 pN accuracy (Figure 1E,F). Ndc80 oligomers attached to nanosprings did not rescue microtubule depolymerization under force (Figure 4). Based on previous observations that microtubule rescues require a dense coating of Ndc80 trimers on the bead surface (Volkov et al., 2018), we conclude that neither a single Ndc80 trimer in presence of Ska, nor a multimer with up to nine Ndc80 copies, is sufficient to rescue depolymerization of the microtubule end. The direct quantification of Ndc80 copy number at the interface between the nanospring and the dynamic microtubule end allows us to conclude that the presence of additional copies of Ndc80, but not Ska, correlates with an increase in the force transmitted by a shortening microtubule.

In summary, we demonstrate the use of the DNA origami nanospring in measuring the force of microtubule motors walking along stable microtubule lattice, as well as the forces exerted by dynamic microtubule ends via proteins following their growth or shortening. In addition, we extend the use of this method to recording single-molecule dynamics of proteins binding and unbinding at the site of force generation. We conclude that the DNA origami nanospring is a powerful tool to study transmission and sensing of microtubule-generated forces *in vitro*.

## Materials and methods

### Design and purification of the DNA nanospring

The originally described DNA nanospring design is based on the M13mp18 (Iwaki et al., 2016), but it could also be adapted to other commercially available scaffolds such as p7560 using caDNAno software (Douglas et al., 2009) and the json file describing the original design.

Step-by-step protocol for modification of the spring design using a different single-stranded DNA scaffold, or to re-staple using a different set of DNA oligos:

- Open the .json file in caDNAno; on the left, in the slice panel, you will see 6 cells are highlighted. This is the cross-section view of potential helices. The right panel (path panel) shows the side view of the DNA construct with cells representing DNA bases. In this design there are only two DNA double strands being formed, 0 and 1 (Figure 1A)
- Zoom in the cells 0 and 1 in the path panel to see DNA strands. The blue line shows the single-strand DNA scaffold which will be folded on itself over cells 0 and 1 via staple DNA oligos which are shown in dark grey.
- Go to the right terminal of scaffold (hold control/command and drag) where you can see the 3’ end (represented as an arrow in caDNAno) and the 5’ end (a square).
- Choose the *select* tool in the control bar in the right and click on 3’ arrow. Once it is selected the arrow turns red. Now you can drag the arrow to extend the scaffold (Supplementary Figure S1A). If there is not enough space to extend the strand go to the rightmost part of the grid and click on the double head arrow. Then you can add more bases to the grid (should be a multiple of 21)
- Repeat the previous step for the 5’ square. Keep in mind that the total length of the blue line should not exceed the length of the actual ssDNA scaffold. Tip: Hold ALT key and click on 3’ or 5’ ends to push them to the extremes of the grid.
- In this design a hairpin of 3 bases is introduced at every 14 bases in one of the scaffold strands (strand 1). Extend this pattern in the newly added bases by clicking on *Insert* in the left bar and then clicking on every 14 squares in the grid (Supplementary Figure S1B).
- Click on hairpins and type the number of bases that you want to have in the hairpin (3). Press Enter to apply. Varying the length of the hairpins will affect the twisting of the resulting spring (Supplementary Figure S1C).
- At this point it is possible to use *Auto Staple* tool on the top bar to generate folding staples and distribute them over the scaffold, but this also resets the present design of staples. Alternatively, you can add staples manually using the *Pencil* tool. Following the existing pattern, insert a complementary oligo next to strands 0 and 1. Connect the oligos by dragging the 3’ of one of them and releasing it over 5’ of the opposite oligo on the other strand of scaffold. This creates a circular oligo that connects two scaffold strands (Supplementary Figure S1D, E).
- Since linear oligos will be used to fold the scaffold, they need to be broken at a certain site. In the current design the breaking site is 11 bases away from the nearest left crossover. Select the Break tool and click on this site to generate the break (Supplementary Figure S1F). *Auto break* tool can be used for the same purpose, but the break sites will be decided by the program and propagated to the whole design, overriding manual settings.
- If you want to use the same staples as for the shorter nanospring, you should count how many bases have been added to the 5’ path of scaffold in caDNAno design. Cut the same number of the bases from 3’ end of the scaffold sequence and paste it before 5’ terminal of the sequence (see Supplement).

Depending on the scaffold used (i.e. M13mp18, p7308, p7560, p7704, p8064, p8100, or p8634) and on the starting position in the scaffold, the same design produces different staple sequences. We provide examples of a json file and caDNAno output for p7560 scaffold in the Supplement.

The scaffold and staple DNA oligos were obtained from Tilibit Nanosystems. Nanospring was folded by mixing 20 nM scaffold and 200 nM staples in the folding buffer (40 mM Tris pH 8.0 with 1 mM EDTA and 12 mM MgCl2) followed by an incubation in a thermocycler: 80°C 10 min, gradient from 80°C to 60°C over 2 hr, gradient from 60°C to 20°C over 2 hr. Folded DNA nanosprings were separated from excess staples and partially folded products on a 1% agarose in Tris-Borate-EDTA buffer supplemented with 12 mM MgCl_2_ (Supplementary Figure S1G). Nanospring-containing bands were excised and extracted from the gel using Freeze ‘N Squeeze™ DNA Gel Extraction Spin Columns.

### Negative stain electron microscopy

A solution of purified nanosprings (3 μL) was placed on a recently glow-discharged grids with a continuous layer of carbon, blotted, and immediately washed three times, followed by the application of 3 μL of the 2% Uranyl acetate solution for 3 min. The grids were then blotted once more with a blotting paper and dried for 20 min. Images were acquired on a JEOL JEM1200 microscope equipped with a TVIPS F416 camera at a nominal magnification of 41000x, resulting in a pixel size of 0.38 nm.

### Protein expression and purification

Dynein motor domain was purified and biotinylated as described (Baclayon et al., 2017; Reck-Peterson et al., 2006). EB3 and MACF2 C-terminus was expressed and purified as described (Rodríguez-García et al., 2020). Ndc80 and Ska were expressed and purified, and Ndc80 was assembled into trimers as described (Huis in ‘t Veld et al., 2019; Volkov et al., 2018).

### Assembly of multimeric Ndc80 modules

Subunits of spyavidin scaffolds were expressed using Traptavidin, Dead-Streptavidin-SpyCatcher, Dead-Streptavidin-SpyTag, and Traptavidin-E6 plasmids. These plasmids are generated by the Howarth laboratory (Chivers et al., 2010; Fairhead et al., 2014) and available through Addgene (26054, 59547, 59548, 59549).

T1S_C_3, T1S_T_3 tetramers and S_C_4 modules were prepared as described previously (Fairhead et al., 2014). To generate multimeric T1S_T_3S_C_12 Spyavidin-scaffolds, T1S_T_3 and S_C_4 tetramers were mixed and purified by size-exclusion chromatography (Superose 6 10/300) as detailed in Figure 4 suppl. panels A and B.

Ndc80 with SPC24^SpyTag^ and SPC25^SortaseHis^ was incubated at 30 μM with T1S_T_3S_C_12 assemblies at 0.3 μM in a 16 hour reaction at 10 °C in Ndc80 buffer. Protease inhibitor (HP PLUS, Serva 39107) was present at 2.5-fold the recommended concentration. 5M Sortase (Hirakawa et al., 2015), Ca^2+^, and GGGGK-TMR peptide (Genscript) were added at concentrations of 10 μM, 10 mM, and 150 μM respectively to fluorescently label Ndc80. The reaction volume was 1 ml and reaction progress was monitored as detailed in Figure 4 suppl. Panel C.

T1S_T_3S_C_12-(Ndc80)x modules were separated from Ndc80 on a 15-50% glycerol gradient of approximately 12 ml in a SW40 rotor (Beckman) for 16 hrs at 4 °C. Manually collected fractions were analyzed as detailed in Figure 4 suppl. Panel D and selected fractions were pooled, frozen in liquid nitrogen, and stored at −80 °C until further use. Shadowing electron microscopy was performed as described previously (Huis in ‘t Veld et al., 2016).

### Coverslip and slide passivation

Glass slides and coverslips were treated with oxygen plasma for 3 min in a PSI Plasma Prep III plasma cleaner at 60 mTorr, 20-50W. Immediately after plasma treatment, the coverslips were immersed into a repel-silane solution (2% dichlorodiethylsilane in trichloroethylene or octamethylcyclooctasilane) for 5 min (Gell et al., 2010; Volkov et al., 2014). After the incubation, silanized coverslips were transfered to 96% ethanol solution and sonicated in a water bath sonicator for 20 min, following by 5-10 rinses with purified water. Silanization was considered successful if the glass was almost dry when emerging from water. Slides and coverslips were dried and stored for up to 2 months.

### Assembly of flow chambers and attachment of nanosprings to the coverslip surface

Silanizing both slides and coverslips provides superior control of non-specific adsorption of proteins to glass but presents a challenge when introducing water solutions into a pre-assembled hydrophobic flow chamber. To overcome this, we used the following sequence (Supplementary Figure S1H). Anti-DIG IgG diluted in in MRB80 (80 mM K-PIPES pH 6.9, 1 mM EGTA, 4 mM MgCl2) to a final concentration of 0.2 μM was placed in 0.5-1μL drops between the strips of double-sided tape (10-15 μL in total) and covered with a piece of silanized coverslip, followed by a 15 min incubation. The chamber was washed with 100 μL MRB80, then with 50 μL 1% Pluronic F-127 in MRB80 and incubated further for 20-60 min. Finally, 10 μL of nanospring diluted in MRB80 was added. Supplementary Figure S1I shows a microscopic field of view after addition of Atto-488 labelled nanosprings diluted 1:10 from a 5 nM stock solution. Alternatively, surface passivation after silanization can be achieved by substituting 1% Pluronic F-127 with 1% tween-20. Tween-20 passivation was shown to be particularly effective against non-specific adsorption of streptavidin (Hua et al., 2014).

### Experiments with dynein

Taxol-stabilized microtubules were prepared by polymerization of 50 μM tubulin (with 5-10% fluorescent-and DIG-labeled tubulin) in MRB80 supplemented with 1 mM GTP and 25% glycerol for 20 min at 37°C, followed by addition of 25 μM taxol and another 10-20 min incubation. Microtubules were sedimented in a Beckman Airfuge at 14 psi for 3 min and resuspended in MRB with 40 μM taxol.

To conjugate nanospring with Qdot and dynein, 4.5 μL nanospring was mixed with 0.5 μL streptavidin-Qdot (f.c. 100 nM) on ice for several hours/overnight. Biotinylated dynein (f.c. 1-20 nM) was mixed with 50 μL MRB80 with 1 mM ATP, 1 mM DTT and 0.5 mg/ml κ-casein, spun in Beckman Airfuge (30 psi, 5 min), and 5 μL of the supernatant was mixed with 5 μL Qdot-nanospring reaction from the previous step.

A flow-chamber prepared with silanized slides and coverslips and passivated with 1% tween-20 as described above was filled with nanospring-Qdot-dynein reaction supplemented with 0.5 mg/ml κ-casein. After washing with MRB80 with 0.5 mg/ml κ-casein, taxol-stabilized microtubules were added and incubated for 3 min, followed by another wash with MRB80 with 0.5 mg/ml κ-casein. Finally, the chamber was filled with Imaging Buffer: MRB80 with 1 mg/ml κ-casein, 50 mM KCl, 1 mM ATP, 40 μM taxol, 20 mM glucose, 4 mM DTT, 0.2 mg/ml catalase, 0.4 mg/ml glucose oxidase.

### Experiments with MACF2

GMPCPP seeds were polymerized by incubating 25 μM tubulin (40% DIG-labelled, total volume 8 μL) and 1 mM GMPCPP for 30 min at 37°C. Polymerized microtubules were sedimented in a Beckman Airfuge (30 psi, 5 min), and the pellet was resuspended on ice in 6 μL MRB80 with addition of 1 mM GMPCPP, followed by a 30 min incubation on ice. The reaction was then transferred to 37°C and microtubules polymerized for 30 min and sedimented as above. The pellet was resuspended in 50 μL MRB80 with 10% glycerol, and aliquots snap-frozen in liquid nitrogen and stored at −80°C for up to 3 months.

Nanosprings were first attached to Qdots as described above. After an incubation on ice, biotinylated MACF2 C-terminus was added at 1 μM to saturate remaining biotin-binding sites on the Qdots. Flow chambers were assembled from silanized slides and coverslips and passivated with 1% tween-20 as described above. Nanospring-Qdot-MACF reaction was then supplemented with 0.5 mg/ml κ-casein and added to the chamber, the chamber was transferred to 37°C. Tubulin polymerization mix containing 15 μM tubulin, 1 mg/ml κ-casein, 0.01% methylcellulose, 1 mM GTP, 20 mM glucose, 4 mM DTT, 0.2 mg/ml catalase, 0.4 mg/ml glucose oxidase, 50 mM KCl, 100 nM EB3 and 15 nM MACF was cleared by centrifugation in a Beckman Airfuge (30 psi, 5 min). Cleared tubulin mix was added to GMPCPP-stabilized seeds and incubated for 10 min at 37°C. Finally, polymerized microtubules were added to the pre-warmed chamber, the chamber was sealed and immediately imaged.

### Experiments with Ndc80 and Ska

Nanospring-Ndc80 trimer conjugation was set up by mixing 10 μL Ndc80 buffer NB (50 mM NaHepes pH 7.5 with 250 mM NaCl and 5% glycerol) with 1 μL nanospring, 0.5 mg/ml κ-casein, 1 mM DTT and 100-200 nM Ndc80 trimers, followed by an incubation for 1-3 hrs on ice. Flow chamber was assembled using silanized slides and coverslips, and passivated with 1% tween-20 as described above. Tubulin polymerization mix was prepared by mixing 1 mg/ml κ-casein, 0.01% methylcellulose, 1 mM GTP, 20 mM glucose, 4 mM DTT, 0.2 mg/ml catalase, 0.4 mg/ml glucose oxidase and 8-10 μM tubulin (5-10% fluorescently labelled) and clearing it in a Beckman Airfuge (30 psi, 5 min). GMPCPP seeds were then added to this mix and incubated at 37°C to start microtubule polymerization (optionally, with addition of 1-100 nM Ska).

The passivated flow-chamber was washed with with 100 μL NB, then with 50 μL NB with 0.5 mg/ml κ-casein and 1 mM DTT. Nanospring-Ndc80 reaction was diluted 3 μL + 7 μL NB with 0.5 mg/ml κ-casein and 1 mM DTT (dilution tuned based on desired nanospring density), and added to the chamber for 3-5 min, followed by a wash with 100 μL MRB80 with 0.5 mg/ml κ-casein and 1 mM DTT. The chamber was then pre-warmed at 37°C, followed by addition of pre-polymerized microtubules and seeds, immediately sealed and imaged at the microscope.

### Imaging and image analysis

Images were acquired using Nikon Ti-E microscope (Nikon, Japan) with the perfect focus system (Nikon) equipped with a Plan Apo 100X 1.45 NA TIRF oil-immersion objective (Nikon), iLas2 ring TIRF module (Roper Scientific) and a Evolve 512 EMCCD camera (Roper Scientific). The sample was illuminated with 488 nm (150 mW), 561 nm (100 mW) and 642 nm (110 mW) lasers through a quad-band filter set containing a ZT405/488/561/640rpc dichroic mirror and a ZET405/488/561/640m emission filter (Chroma). Images were acquired sequentially with MetaMorph 7.8 software (Molecular Devices, San Jose, CA). The final resolution was 0.107 μm/pixel, using an additional 1.5x lens. The objective was heated to 34°C by a custom-made collar coupled with a thermostat, resulting in the flow chamber being heated to 30°C. TIRF penetration depth was fine-tuned separately for each fluorescent channel.

Further analysis has been done in Fiji (Schindelin et al., 2012) and Julia using custom scripts available at https://github.com/volkovdelft/kymo.jl. Kymographs were made through a reslice operation using the kymograph_3channel.ijm macro. Position of the nanospring end was determined by running kymoNS.ipynb in a jupyter notebook and following in-line comments. In brief, the script opens the kymograph and waits for the user’s clicks to 1) select a portion of the kymograph with only one particle to trace, 2) select a ‘background’ region and the initial position of the particle. Then each line of the kymograph is fit to a gaussian to determine localization of a particle and its brightness.

### Preparation of beads and optical trapping

Glass 1 μm beads were covalently bound to poly-L-lysine grafted with biotinylated poly-ethyleneglycole (PLL-PEG) as described previously (Volkov et al., 2018). Biotins on the bead surface were then saturated with neutravidin. Optical trapping was performed using a custom instrument described before (Baclayon et al., 2017). To calibrate the nanospring force-extension curve, neutravidin-coated beads were introduced into a flow-chamber with DIG-nanospring-biotin bound to the surface through anti-DIG IgG (see above). After a bead was trapped, the piezo stage was moved manually in 100-nm steps to scan the bead displacements around the nanospring attachment point. The beads that did not produce a bell-shaped displacement profile, like the one shown in Figure 1D, were discarded from further analysis.

For experiments with dynein, biotinylated dynein motor domains were bound to streptavidin-coated PLL-PEG beads. We tuned surface density of dynein such that 30% or lower of beads interacted with a microtubule, ensuring predominantly single-motor events. Experiments were performed in a buffer that contained MRB80 with 1 mg/ml κ-casein, 50 mM KCl, 1 mM ATP, 40 μM taxol, 20 mM glucose, 4 mM DTT, 0.2 mg/ml catalase, 0.4 mg/ml glucose oxidase. Position of the bead was recorded with a quadrant photo detector at 10 kHz, and simultaneously using differential interference microscopy to monitor bead-microtubule interaction. Experiments with MACF-coated beads were performed as described previously (Rodríguez-García et al., 2020).

## Supporting information

nanospring_caDNAno.json

## Acknowledgements

We are grateful to Jean-Philippe Sobczak (Tilibit Nanosystems) for discussions and assistance with design of the modified nanosprings, to Ruben Kazner and Isabelle Stender (MPI Dortmund) for assistance preparing Ndc80 multimers, and to Nemo Andrea (TU Delft) for help with negative stain EM. We also thank Marian Baclayon and Esengül Yildirim (TU Delft) for dynein purification, and all members of the Dogterom lab for discussions. This work was supported by an EMBO short-term fellowship (grant 7203) to PJH and European Research Council Synergy Grant MODELCELL (grant 609822) to MD and AA.

---

**Supplementary Figure S1.**
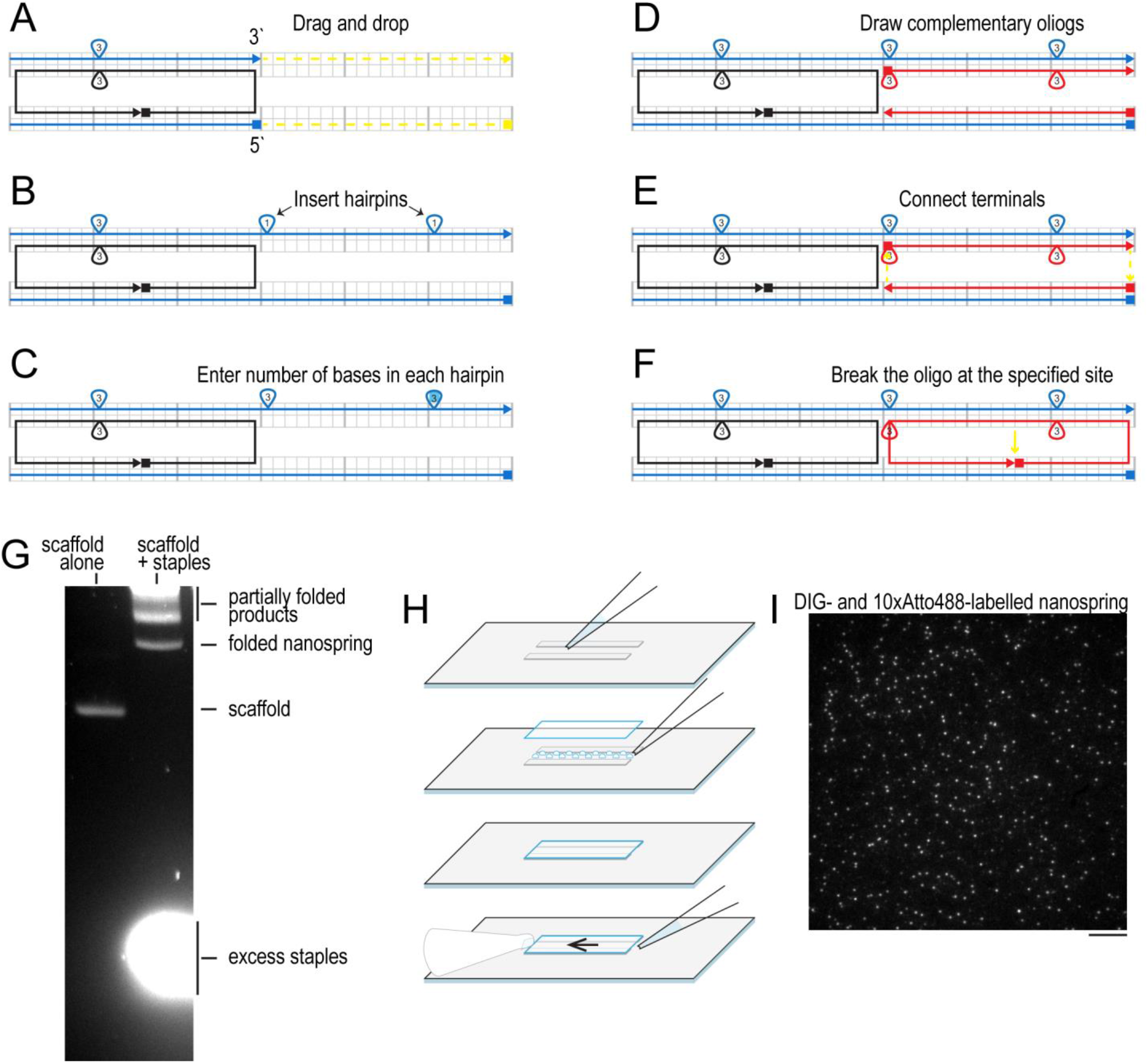
(A-F) step by step visual guide to extend an existing nanospring design using caDNAno. (G) Purification of nanosprings by agarose gel electrophoresis, shown are lanes with unfolded scaffold in absence of staples (left), and the products of folding (right). (H) Schematic diagram showing assembly of a flow chamber using silanized slides and coverlsips. (I) Typical microscopic field of view with 10x Atto-labelled nanosprings attached to glass surface using DIG and anti-DIG IgG. Scale bar: 10 μm.

**Suppelementary Figure S2.**
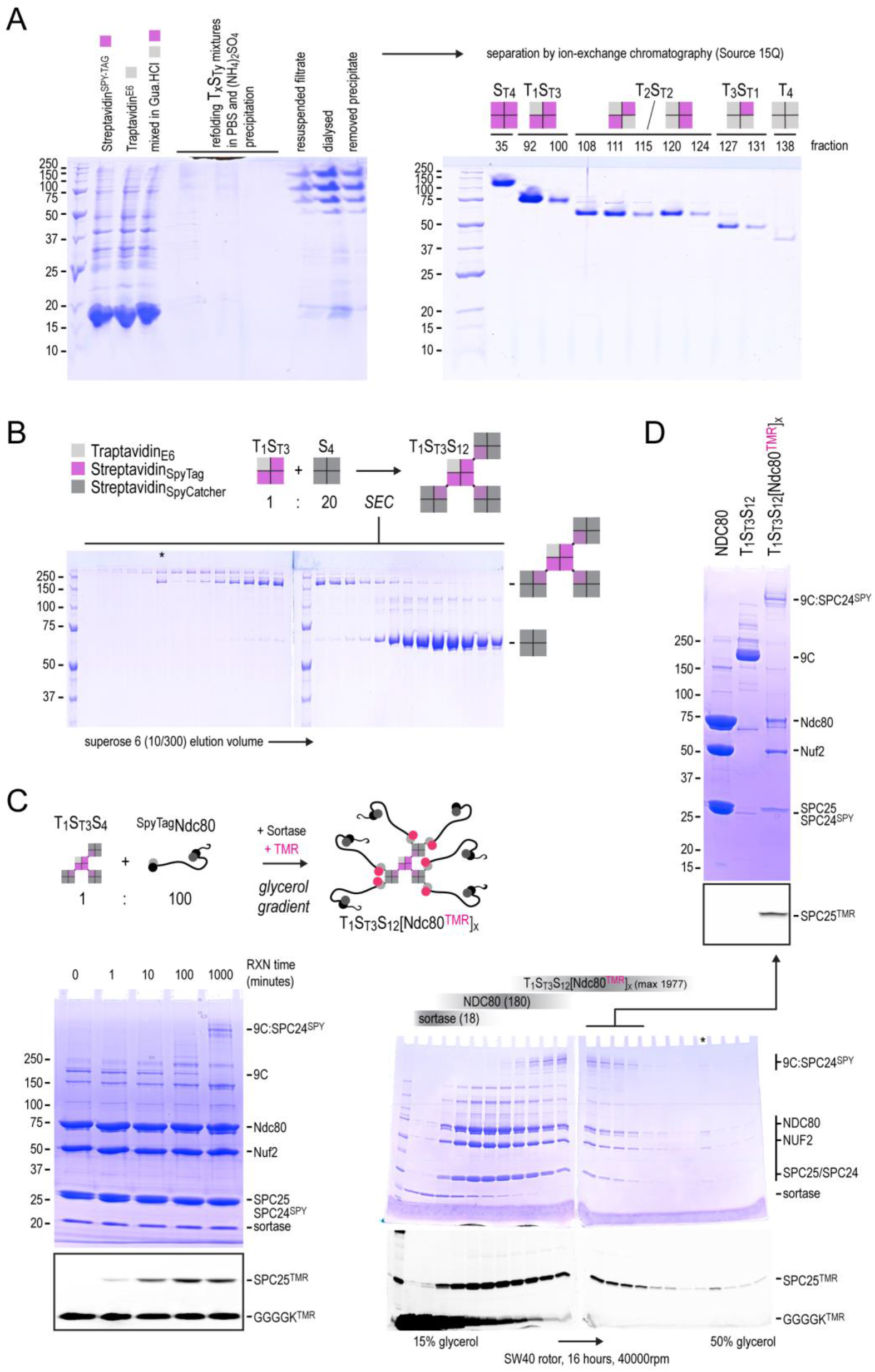
(A) Monomeric Traptavidin and Streptavidin-SpyTag were mixed, refolded into tetramers in PBS, and separated using ion-exchange chromatography as described (Fairhead et al., 2014). Streptavidin remains a tetramer during SDS-PAGE since samples were not heated before analysis. (B) T1S_T_3S_C_12 scaffolds were generated from T1S_T_3 and S_C_4 tetramers and purified by size-exclusion chromatography (Superose 6 10/300). Fractions containing T1S_T_3S_C_12 were pooled and stored for further use. S_C_ subunits are shown in grey when the SpyCatcher module is free to react and in light purple when covalently bound to S_T_ subunits. The asterisk indicates material from a later eluting fraction that was loaded by mistake. Streptavidin remains multimeric during SDS-PAGE since samples were not heated before analysis. (C) A large molar excess of Ndc80 was bound to the T1S_T_3S_C_12 scaffolds and simultaneously fluorescently labeled. Streptavidin and Streptavidin-SPC24 assemblies remain multimeric during SDS-PAGE since samples were not heated before analysis. In-gel fluorescence analysis and coomassie staining of the same gels. (D) SDS-PAGE analysis of the reaction mixture separated on a glycerol density gradient. Peptide, sortase, monomeric Ndc80, and multimeric Ndc80 are separated into different zones of the gradient. Selected fractions containing T1S_τ_3S_C_12-(Ndc80)x modules are indicated. Streptavidin and Streptavidin-SPC24 assemblies remain multimeric during SDS-PAGE since samples were not heated before analysis. In-gel fluorescence analysis and Coomassie staining of the same gels. The final product is also shown side-by-side with monomeric Ndc80 and the spyavidin scaffold on a separate gel.

